## Supplemental Information for "Clonal associations of lymphocyte subsets and functional states revealed by single cell antigen receptor profiling of T and B cells in rheumatoid arthritis synovium"

**Table S1.** Clinical metadata for patient cohort.
**Table S2.** Differentially expressed genes among clusters for examined subsets.
**Table S3.** Gene signatures utilized in this study.
**Table S4.** Antibodies utilized in this study.

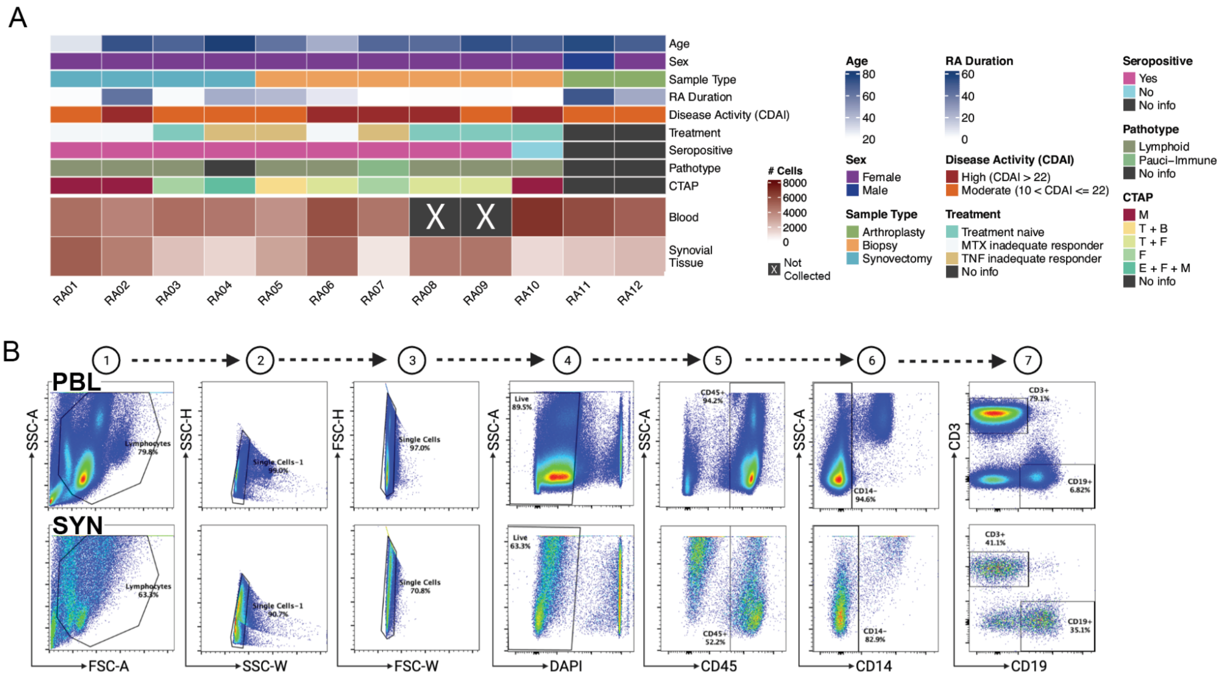


**Figure S1. A.** Heat map highlighting the cell recovery for each patient, along with metadata associated with the cohort. **B.** Step-wise flow sorting scheme for blood and synovial tissue samples.

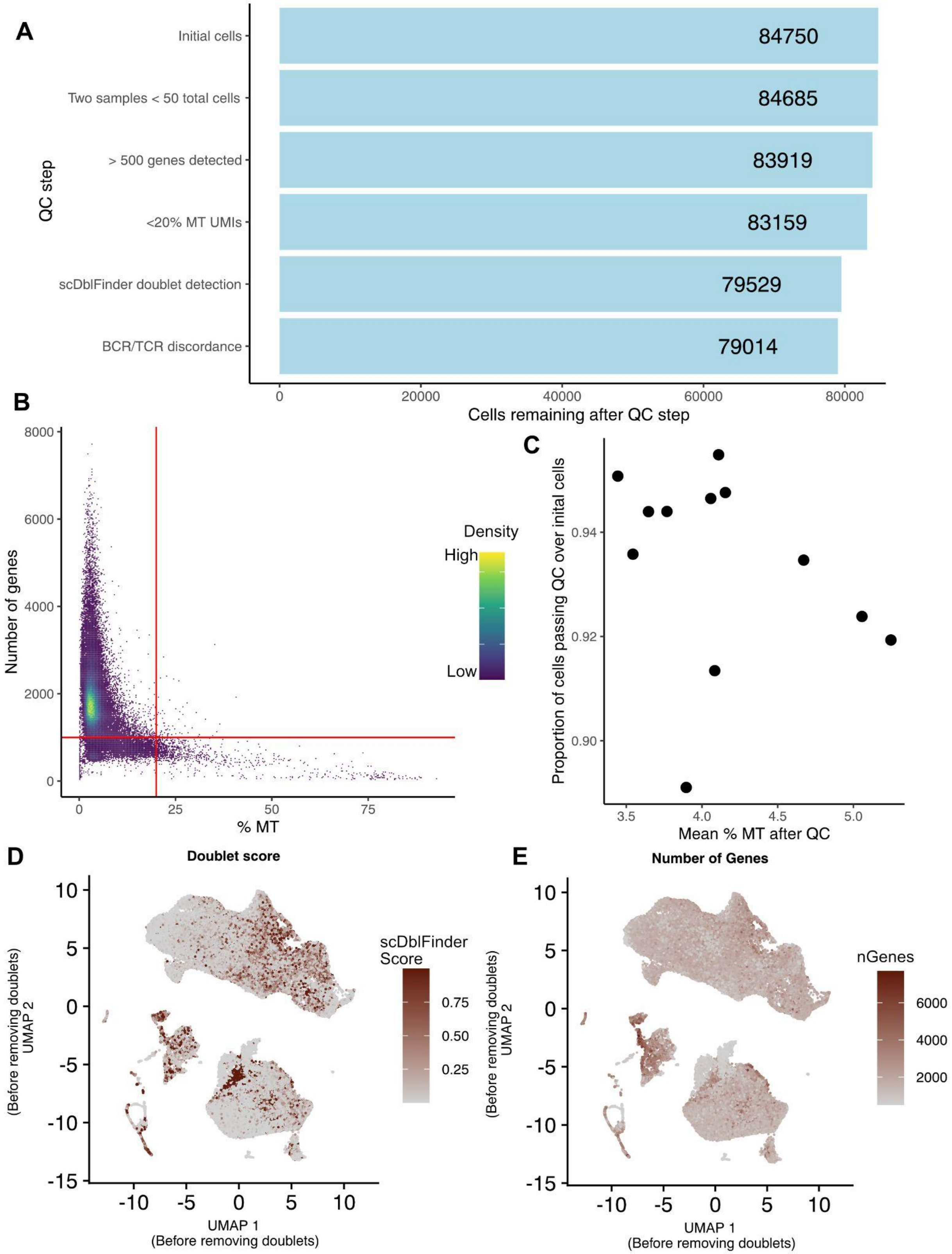

A
QC step
Initial cells
Two samples < 50 total cells
> 500 genes detected
<20% MT UMIs
scDblFinder doublet detection
BCR/TCR discordance
84750
84685
83919
83159
79529
79014
Cells remaining after QC step
B
Number of genes
% MT
Density
High
Low
C
Proportion of cells passing QC over inital cells
Mean % MT after QC
D
Doublet score
scDblFinder Score
UMAP 1 (Before removing doublets)
UMAP 2 (Before removing doublets)
E
Number of Genes
nGenes

**Figure S2. A.** Bar plot of initial quality control filters showing the number of cells remaining after each quality control step. **B.** Scatter plot of the number of detected genes and the percent of mitochondrial reads for each cell. Red lines correspond to the selected QC filters (>500 genes and <20% MT reads). **C.** Scatter plot of mean mitochondrial percentage and proportion of cells passing QC for each sample. Each dot corresponds to a single sample. **D.** UMAP showing doublet scores according to scDblFinder of all cells before QC. **E.** UMAP showing the number of genes in each cell before QC.

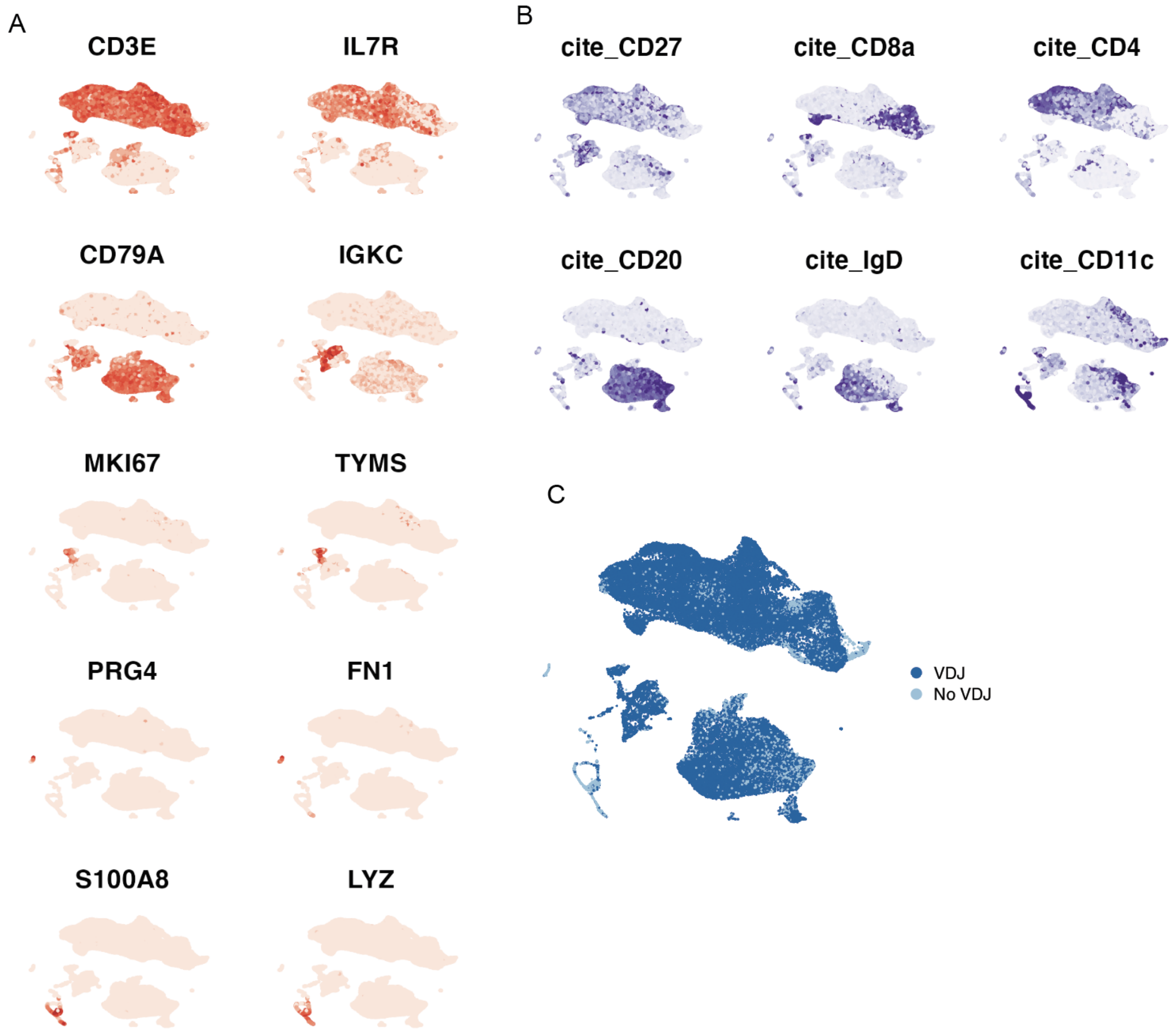


**Figure S3. A.** UMAPs of the gene expression levels for select markers used to identify each population. **B.** UMAPs of the detection of CITEseq barcoded antibodies used in this study. **C.** UMAP of the distribution of cells that have associated VDJ information.

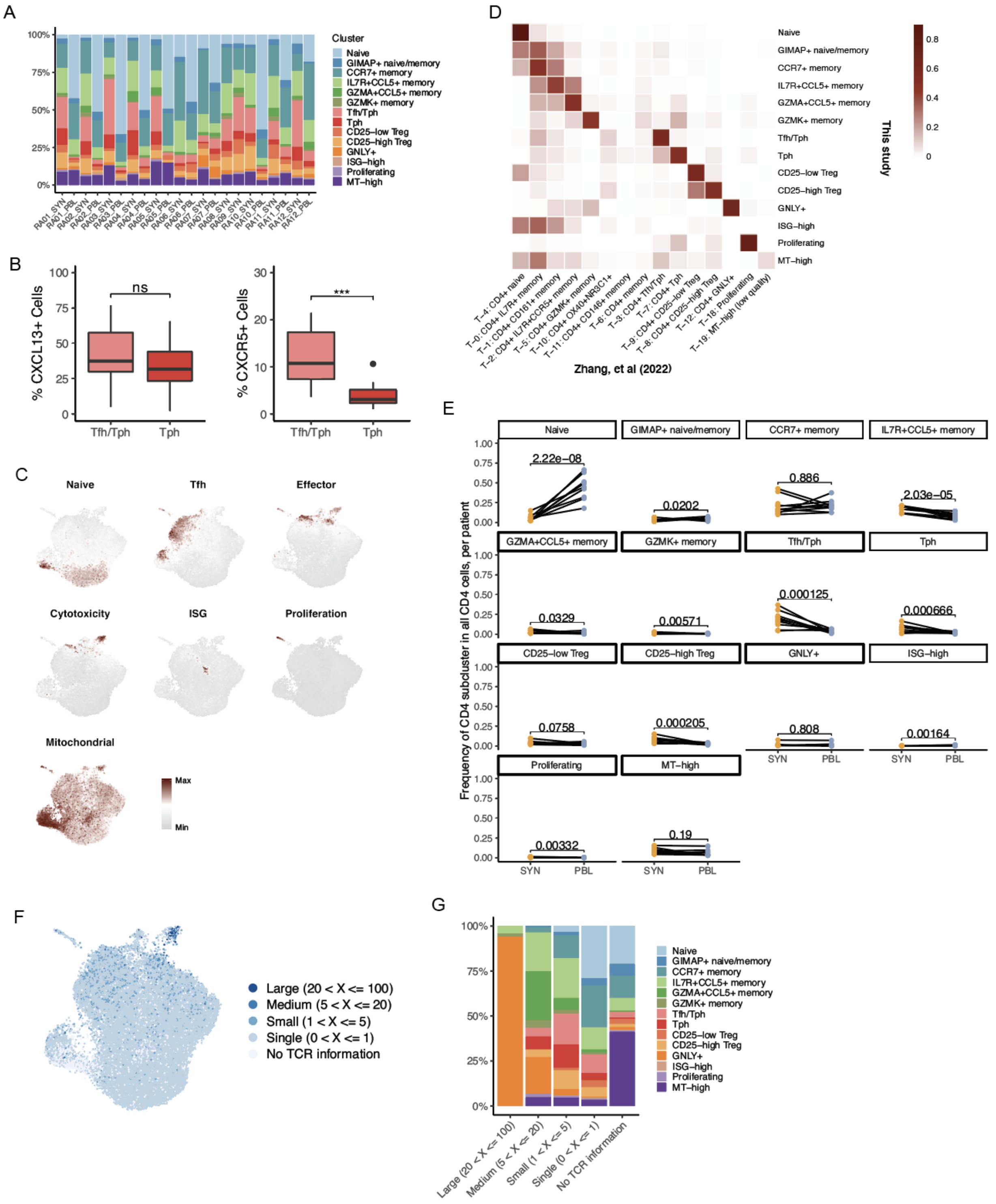


**Figure S4. A.** Bar plot of the CD4+ cluster composition for each sample. **B.** Box plots of the percent of cells per patient expressing *CXCL13* and *CXCR5* in the Tfh/Tph and Tph clusters. **C.** UMAPs highlighting the enrichment of select gene signatures. **D.** Heat map of the confidence for each cluster to match the CD4+ T cell clusters of a previously-described single-cell synovial tissue dataset. **E.** For each cluster, frequency of cell representation as a proportion of all CD4 cells in a sample. Each dot represents a single sample, and lines denote paired blood and tissue for a donor. Significance is determined by paired T-tests, with multiple testing correction. **F.** UMAP of the clonal expansion among CD4+ cells. **G.** Bar plot of the distribution of clusters across each clone size category. *** $p \leq 0.001$; ns, not significant.

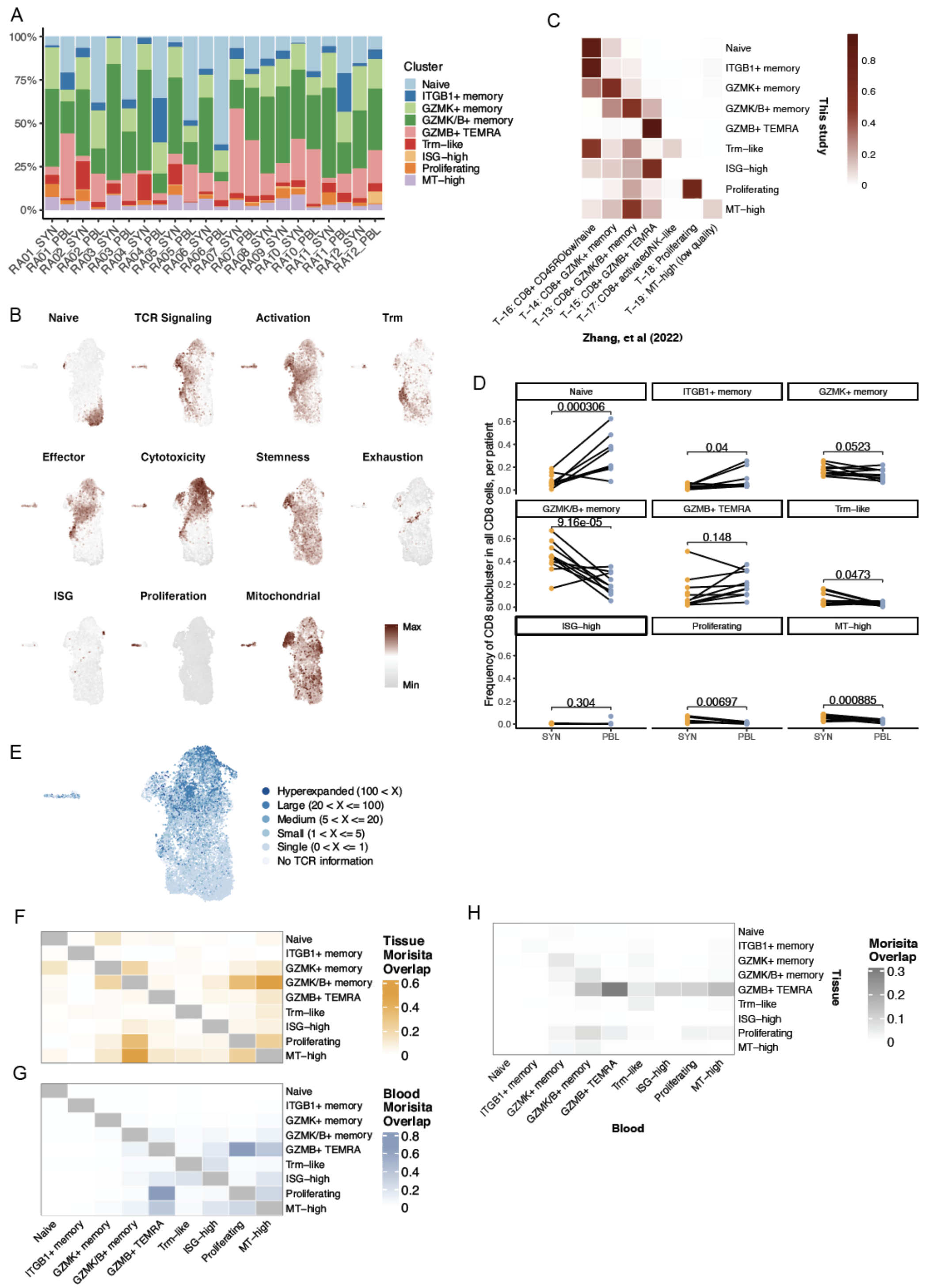

**Figure S5. A.** Bar plot of the CD8+ cluster composition for each sample. **B.** UMAPs highlighting the enrichment of select gene signatures. **C.** Heat map of the confidence for each cluster to match the CD8+ T cell clusters of a previously-described single-cell synovial tissue dataset. **D.** For each cluster, frequency of cell representation as a proportion of all CD4 cells in a sample. Each dot represents a single sample, and lines denote paired blood and tissue for a donor. Significance is determined by paired T-tests, with multiple testing correction. **E.** UMAP of the clonal expansion among CD8+ cells. **F and G**. Heat map of pairwise clonal overlap values calculated using Morisita's index for synovial tissue **(F)** and blood **(G)**. **H.** Heat map of pairwise clonal overlap values between tissue and blood calculated using Morisita's index.

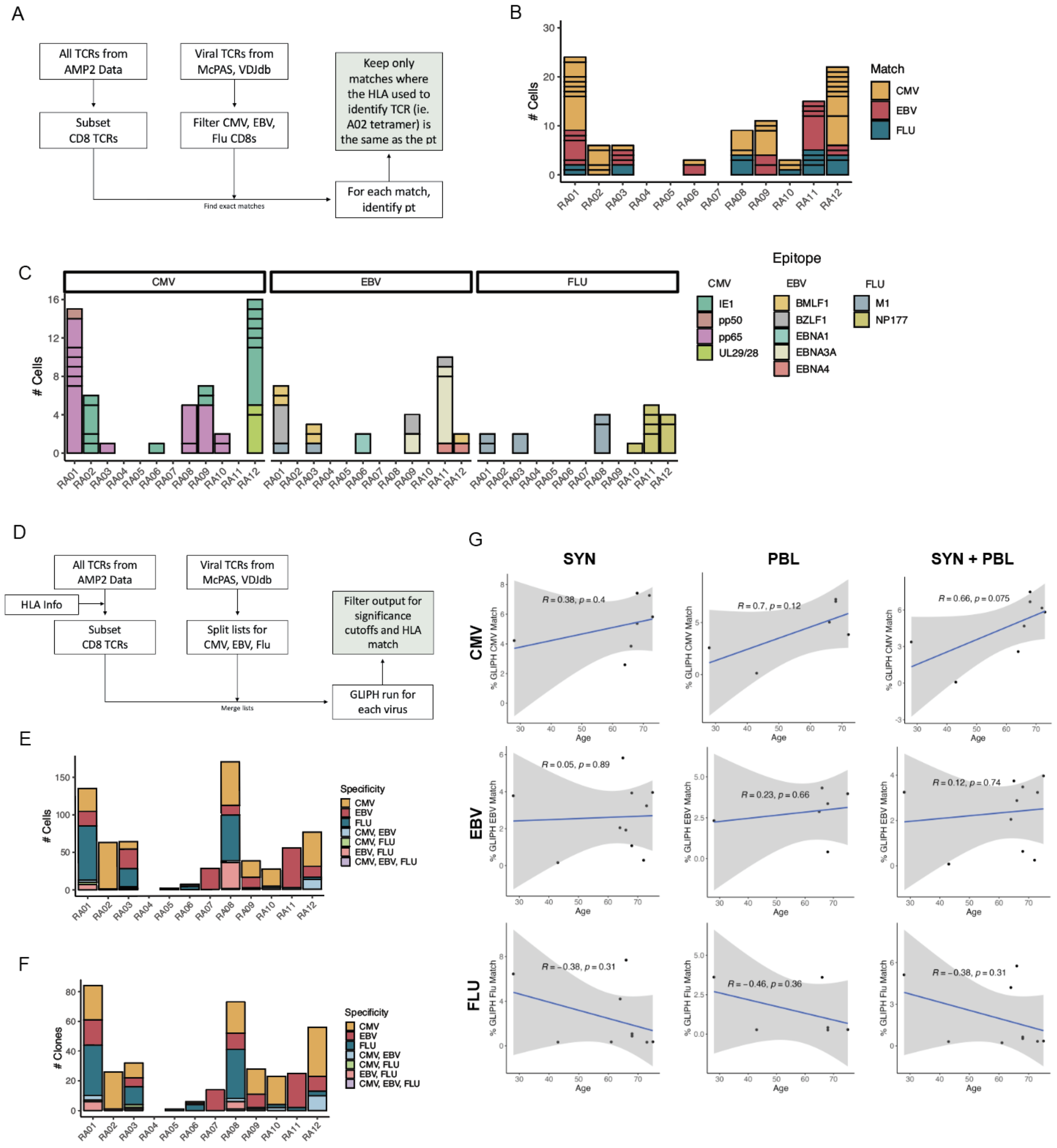

**Figure S6. A.** Schematic diagram of exact TCR matching with viral-specific databases. **B.** Bar plot of the number of exact viral-reactive matching cells per patient, split by virus. Box size denotes size of clone. **C.** Bar plot of exact matches per patient, split by virus and epitope. **D.** Schematic diagram of TCR motif finding with viral-specific databases using GLIPH2. **E.** Bar plot of the number of GLIPH2 motif viral-reactive matching cells per patient, split by virus. **F.** Bar plot of unique clones with virus match using GLIPH2 per patient, split by virus. **G.** Scatter plots of age and percent virus matching CD8+ T cells using GLIPH2, split by tissue.

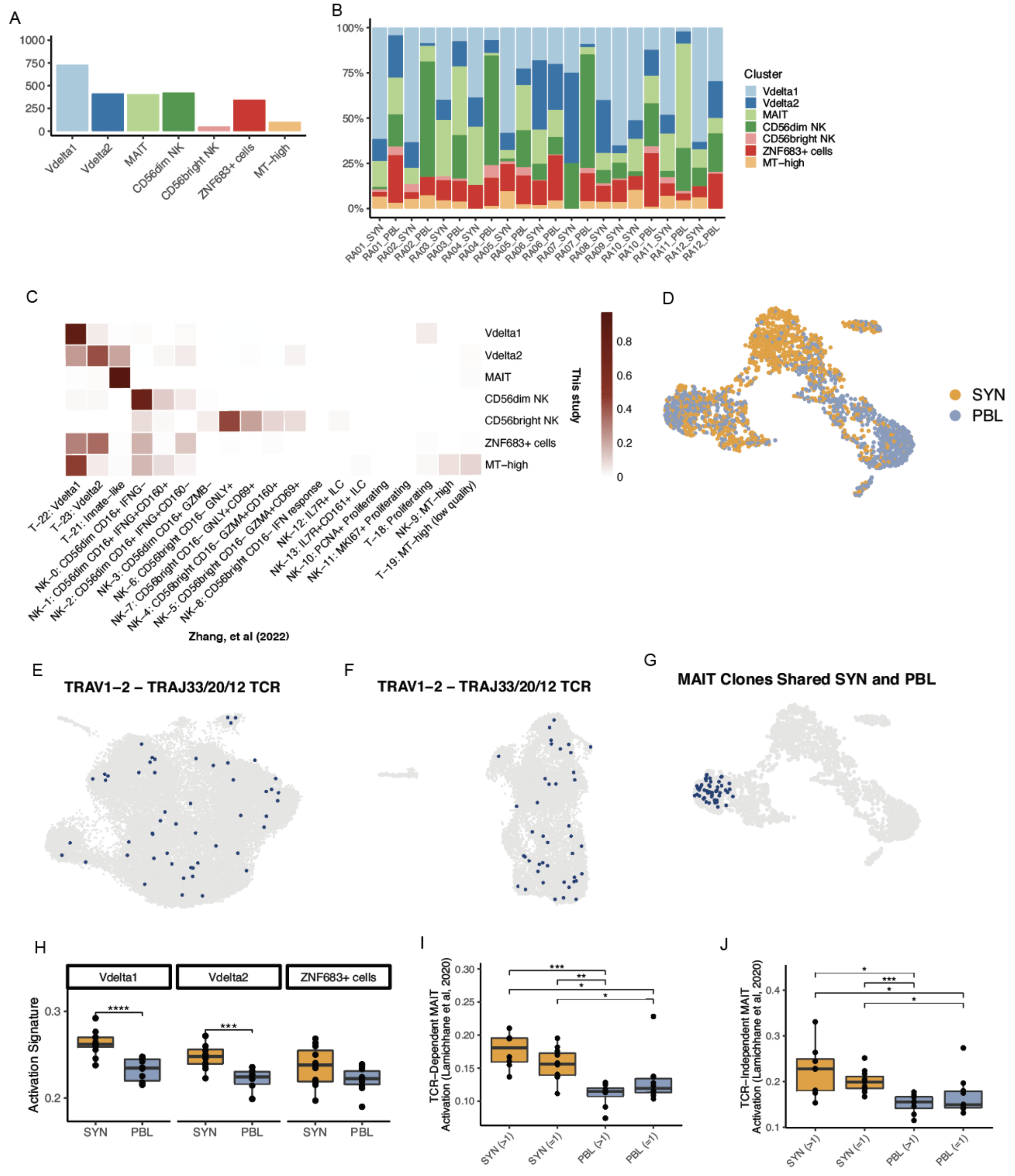

**Figure S7. A.** Bar plot of the number of cells included in each cluster. **B.** Bar plot of the CD4+ cluster composition for each sample. **C.** Heat map of the confidence for each cluster to match the innate T and NK cell clusters of a previously-described single-cell synovial tissue dataset. **D.** UMAP showing the distribution of tissue source. UMAPs highlighting CD4+ **(E)** and CD8+ **(F)** T cells with a TRAV1-2 and either TRAJ33, 20, or 12 gene rearrangement. **G.** UMAP highlighting MAIT clones found in both blood and synovial tissue of a donor. **H.** Box plot of activation signature across innate T cell clusters. **I and J.** TCR-dependent **(I)** and independent **(J)** MAIT cell activation signatures from Lamichhane et al, 2020, split by tissue and clonal expansion.

Significance calculated using paired Wilcox testing with Holm correction. * $p \le 0.05$; ** $p \le 0.01$; *** $p \le 0.001$; **** $p \le 0.0001$; ns, not significant.

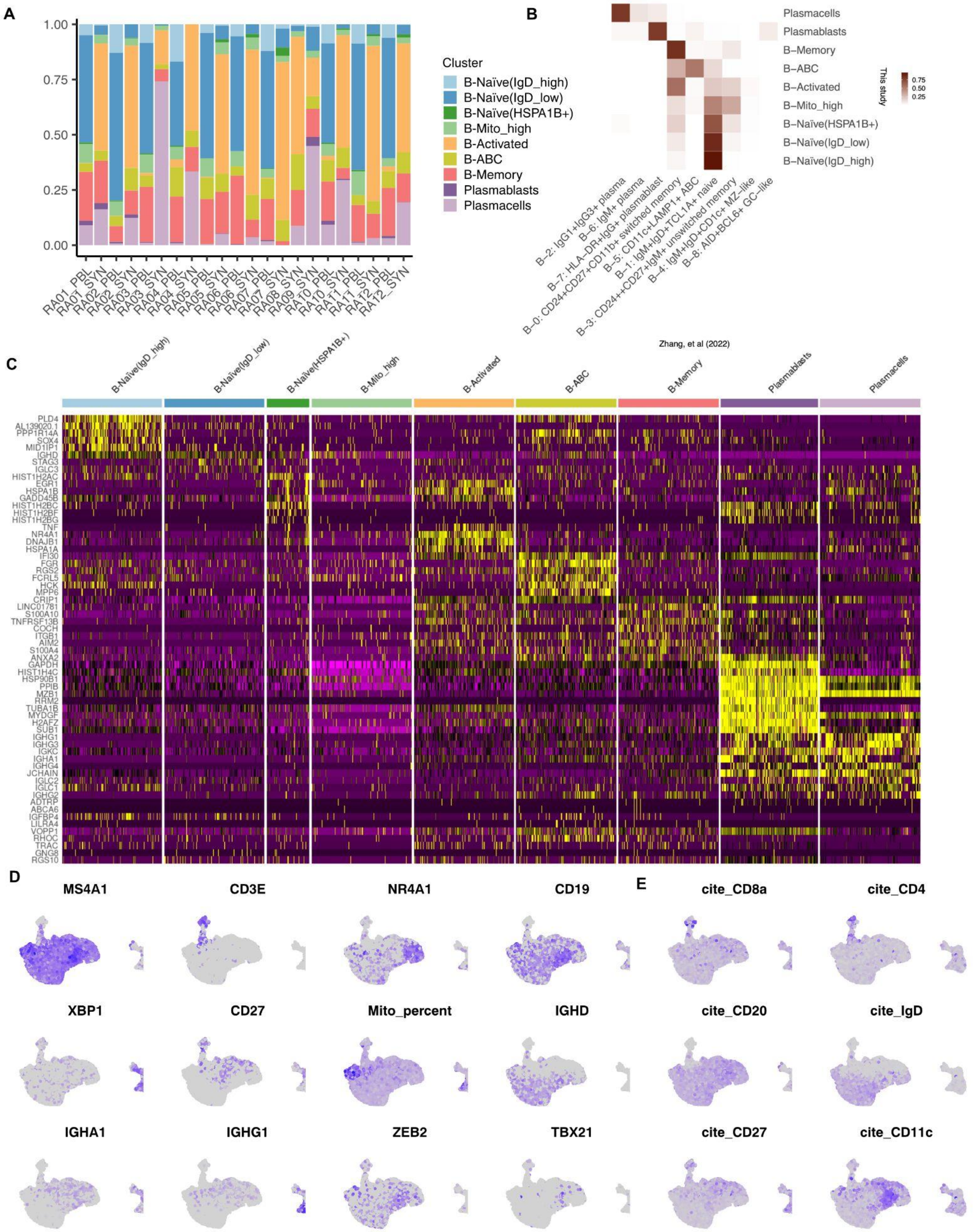
A
B
C
D
E
Cluster
B-Naïve(IgD_high)
B-Naïve(IgD_low)
B-Naïve(HSPA1B+)
B-Mito_high
B-Activated
B-ABC
B-Memory
Plasmablasts
Plasmacells
1.00
0.75
0.50
0.25
0.00
RA01_PBL
RA01_SYN
RA02_PBL
RA02_SYN
RA03_PBL
RA03_SYN
RA04_PBL
RA04_SYN
RA05_PBL
RA05_SYN
RA06_PBL
RA06_SYN
RA07_PBL
RA07_SYN
RA08_SYN
RA09_SYN
RA10_PBL
RA10_SYN
RA11_PBL
RA11_SYN
RA12_PBL
RA12_SYN
Plasmacells
Plasmablasts
B−Memory
B−ABC
B−Activated
B−Mito_high
B−Naïve(HSPA1B+)
B−Naïve(IgD_low)
B−Naïve(IgD_high)
This study
0.75
0.50
0.25
B−2: IgG1+IgG3+ plasma
B−6: IgM+ plasma
B−7: HLA−DR+IgG+ plasmablast
B−0: CD24+CD27+CD11b+ switched memory
B−5: CD11c+LAMP1+ ABC
B−1: IgM+IgD+TCL1A+ naive
B−3: CD24++CD27+IgM+ unswitched memory
B−4: IgM+IgD+CD1c+ MZ−like
B−8: AID+BCL6+ GC−like
Zhang, et al (2022)
B-Naïve(IgD_high)
B-Naïve(IgD_low)
B-Naïve(HSPA1B+)
B-Mito_high
B-Activated
B-ABC
B-Memory
Plasmablasts
Plasmacells
PLD4
AL139020.1
PPP1R14A
SOX4
MID1IP1
IGHD
STAG3
IGLC3
HIST1H2AC
EGR1
HSPA1B
GADD45B
HIST1H2BC
HIST1H2BF
HIST1H2BG
TNF
NR4A1
DNAJB1
HSPA1A
IFI30
FGR
RGS2
FCRL5
HCK
MPP6
CRIP1
LINC01781
S100A10
TNFRSF13B
COCH
ITGB1
AIM2
S100A4
ANXA2
GAPDH
HIST1H4C
HSP90B1
PPIB
MZB1
RRM2
TUBA1B
MYDGF
H2AFZ
SUB1
IGHG1
IGHG3
IGKC
IGHA1
IGHG4
JCHAIN
IGLC2
IGLC1
IGHG2
ADTRP
ABCA6
IGFBP4
LILRA4
VOPP1
RHOC
TRAC
GNG8
RGS10
MS4A1
CD3E
NR4A1
CD19
XBP1
CD27
Mito_percent
IGHD
IGHA1
IGHG1
ZEB2
TBX21
cite_CD8a
cite_CD4
cite_CD20
cite_IgD
cite_CD27
cite_CD11c

**Figure S8. A.** Bar plot of the B cell cluster composition for each sample. **B.** Heat map of the confidence for each cluster to match the B cell clusters of a previously described single-cell synovial tissue dataset. **C.** Heatmap of top 10 differentially expressed genes for each cell population according to average log2-fold change. **D.** UMAP of additional markers used to identify cell types. **E.** UMAP of cite seq markers available in this dataset.

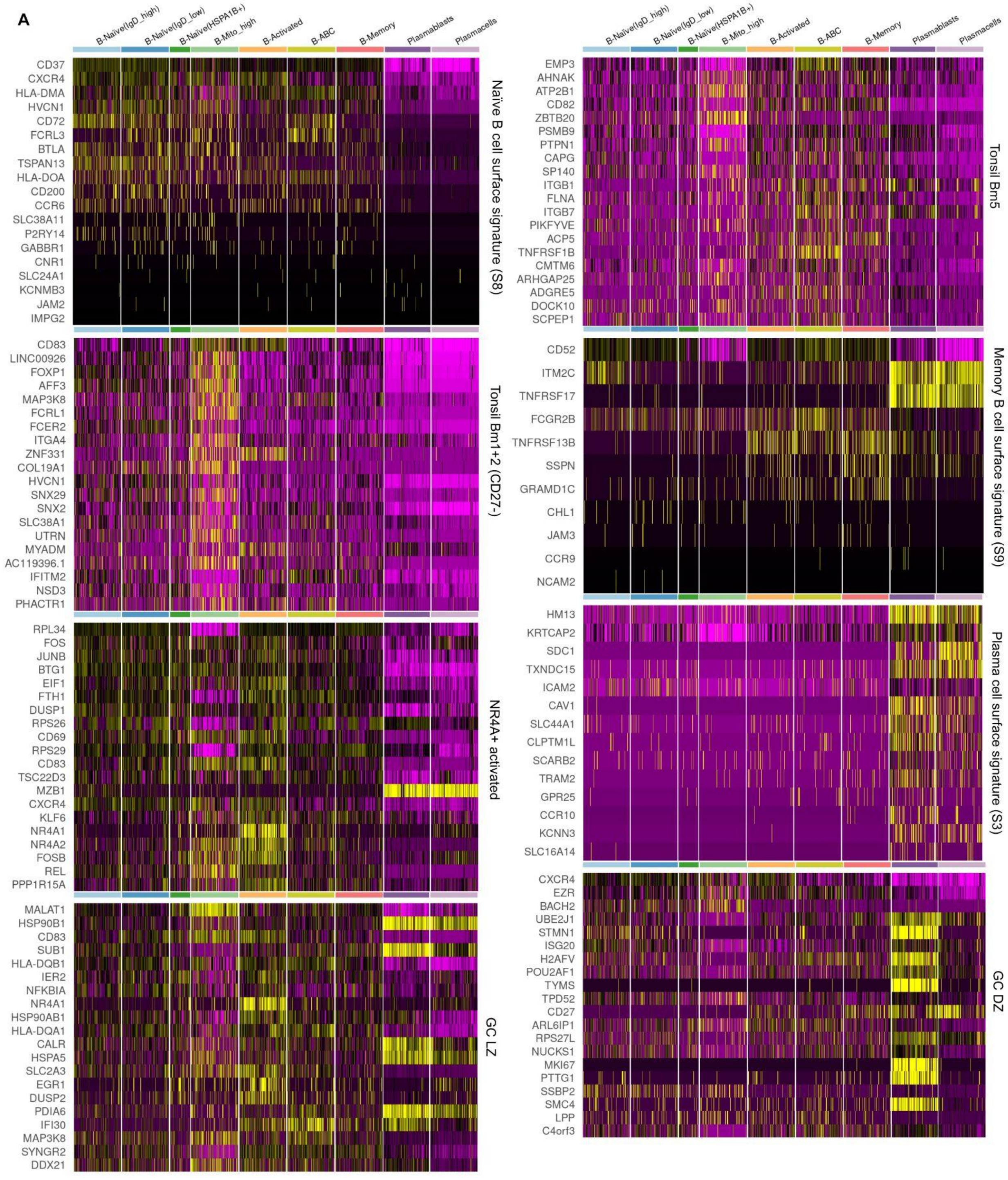


**Figure S9. A.** Heatmaps of 8 selected gene signatures from Figure 5E.

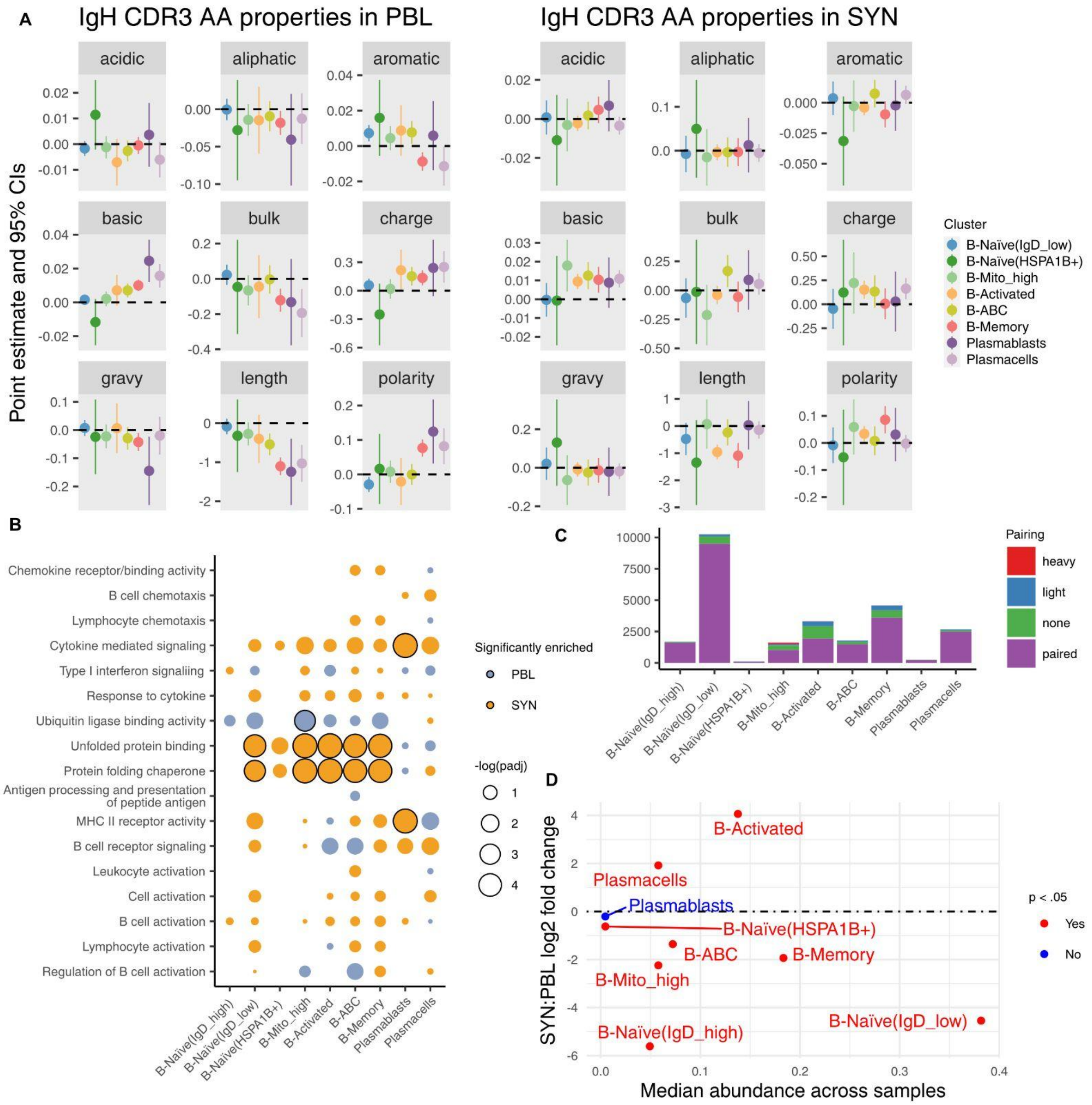


**Figure S10. A.** Plot showing the effect size and 95% confidence intervals of population on IgH CDR3 amino acid properties using all B-Naive(IgD-high) cells (SYN and PBL) as the reference population, split by tissue. **B.** Bar chart of Immunoglobulin chain pairings for each cell population. **C.** Dotplot of selected pathways checked for gene set enrichment between blood and synovium cells for each population. Dots outlined in black deemed significantly enriched. p.adjust < .05. **D.** Plot showing the median sample abundance of each cluster and the synovium to peripheral blood log2 fold change for each cluster, significance determined according to mixed-effect model specified using MASC.

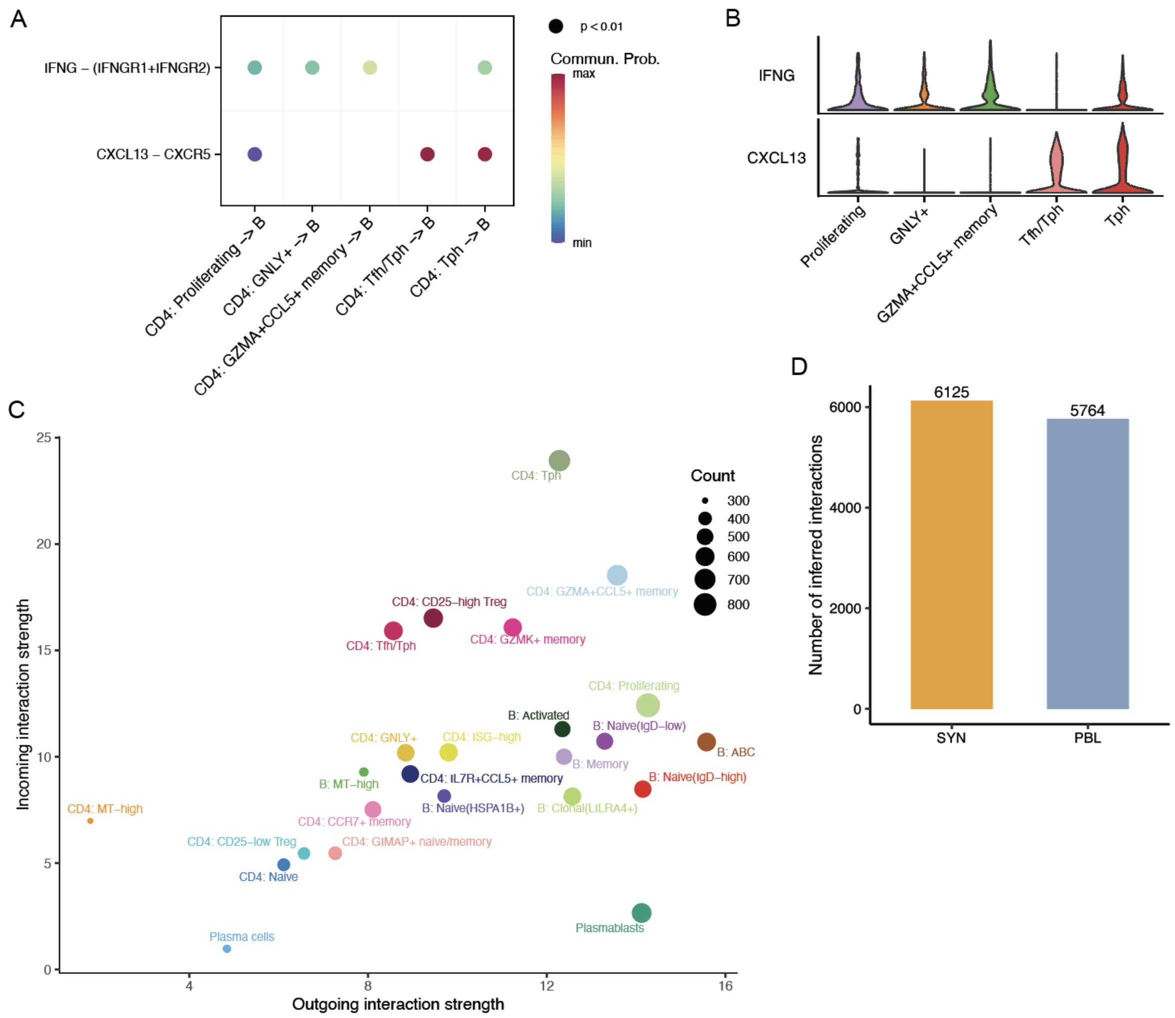


**Figure S11. A.** Dot plot of select signaling pathways between CD4+ T cell subsets and B cells. **B.** Violins of the expression of IFNG and CXCL13 in the CD4+ T cell subsets identified in (A). **C.** Scatter plot of the computed incoming and outgoing signaling strengths for each CD4+ T and B cell subpopulation. **D.** The number of significant interactions detected in each tissue.
